# Brain network dynamics in transitions of consciousness reorganize according to task engagement

**DOI:** 10.1101/2023.06.08.544178

**Authors:** Samika S. Kumar, Anat Arzi, Corinne Bareham, Javier Gonzalez-Castillo, Isabel Fernandez, Enzo Tagliazucchi, Pedro A.M. Mediano, Peter A. Bandettini, Tristan A. Bekinschtein

**Author notes:** Corresponding author: Samika Kumar. Declaration of interests The authors declare no competing interests.

## Abstract

Substantial changes in behavior, physiology, and brain function occur when alertness decreases ^1– 5^. These changes in brain function involve increased synchronization between cortical areas ^6,7^ as well as alterations in sensory processing pathways and networks connecting the thalamus and cortex ^5,8–11^. Cognitive tasks engage overlapping functional networks with sensory pathways facilitating information processing ^12,13^, and thalamocortical and corticocortical networks supporting task performance ^14,15^. Frontoparietal circuits play a crucial role in cognitive tasks ^16^ and states of decreased consciousness ^17^. To develop an integrated framework of consciousness and cognition, it is important to understand how fluctuations in alertness and cognitive processing interact in these shared circuits ^18^. Our hypothesis is that during periods of low alertness, individuals who actively maintain task engagement would recruit additional frontoparietal and sensory processing networks, while thalamocortical dynamics that typically change during sleep onset would remain unaffected. Our findings demonstrated that as alertness decreased, passively listening to auditory tones led to increased synchronization in the parietal lobe, whereas actively performing an auditory task resulted in increased long-range frontoparietal synchronization. During decreasing alertness, passive listening (but not active task engagement) was associated with widespread increased synchronization between the thalamus and cortex. In contrast, active task engagement (but not passive listening) led to increased synchronization between the auditory cortex and the rest of the brain. These results reveal the functional mechanisms of the brain’s flexible reorganization during transitions of consciousness when individuals are actively engaged in cognitive processes.

## RESULTS

We characterized how auditory, thalamocortical, and frontoparietal networks change during decreasing alertness across two different levels of task engagement (pre-registered https://osf.io/nua4y) (**Figure 1a**). We identified changes that were 1) common to both passive listening and active task engagement, 2) specific to active task engagement, and 3) specific to passive listening (**Figure 1b**). Healthy adult participants listened to binaural auditory tones during simultaneously acquired electroencephalography (EEG) and functional magnetic resonance imaging (fMRI) (EEG-fMRI). In an *Active* task, participants (N=20) indicated whether each tone was heard to the left or right of their midline using a button box, and in a *Passive* task, participants (N=20) passively listened to the tones with no motor response required (**Figure 1c**). We extracted the fMRI timeseries for 188 cortical brain regions of interest (ROIs) grouped into six networks (i.e., Visual, Auditory, Somatomotor, Dorsal Attention, Ventral Attention, Control, and Default Mode Network (DMN)), and two ROIs for the left and right thalamus ^19,20^. At each timepoint, instantaneous phase coherence was calculated between each pair of ROI timeseries, resulting in one edge timecourse for each ROI pair ^21^ (**Figure 1d**). The concurrent EEG data were classified as alert or drowsy (i.e., high or low levels of alertness) on four-second epochs ^22^ (**Figure 1e**).

**Figure 1:**
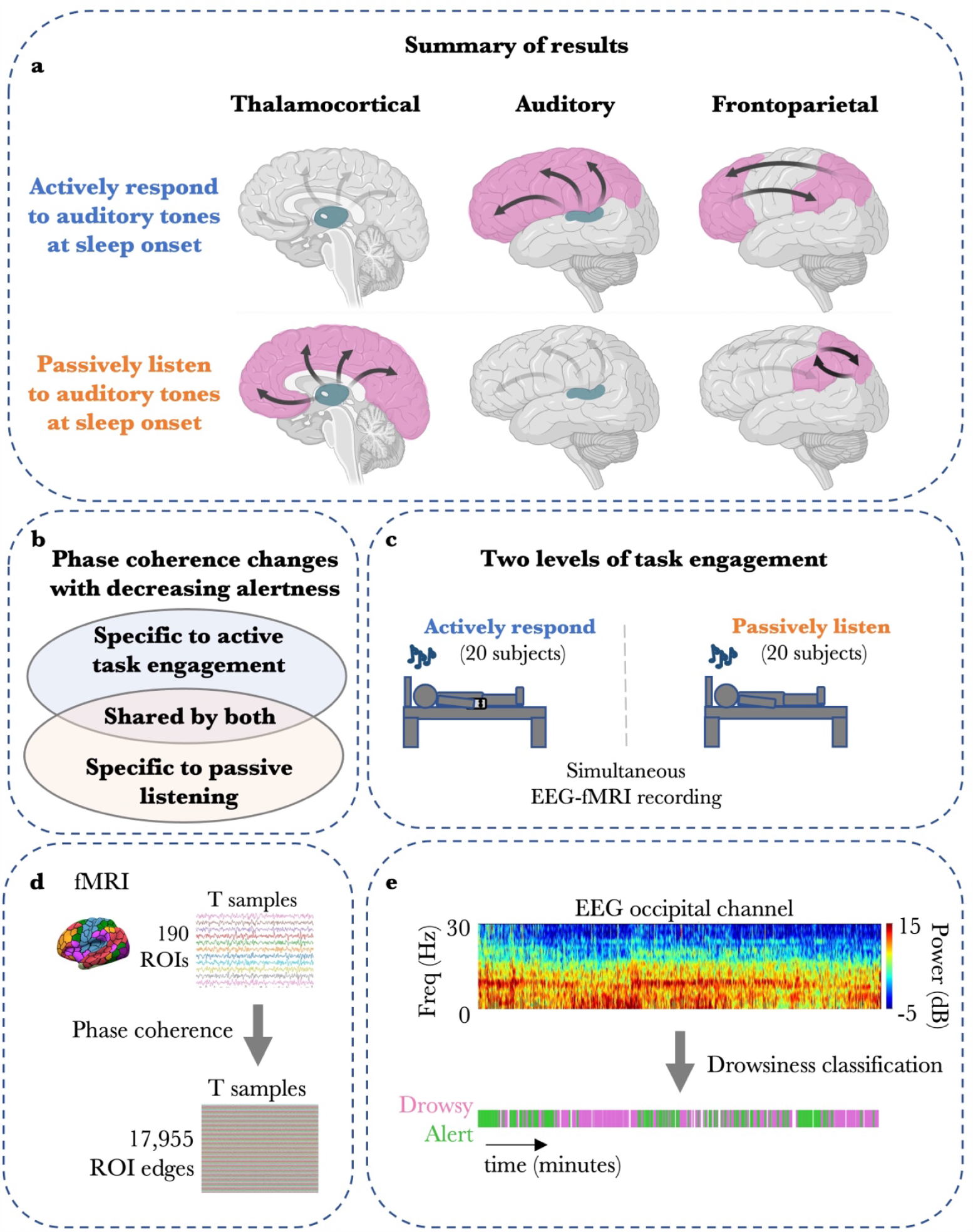
Summary, experimental design, and analysis. **a**. Summary of main findings. At sleep onset, the *Active* task is associated with relatively unchanged thalamocortical dynamics, as well as increased auditory-cortical and frontoparietal synchronization. In contrast, the *Passive* task is associated with increased thalamocortical and parietal synchronization, and auditory-cortical dynamics are relatively unchanged. **b**. We explored changes in fMRI phase coherence that are specific to each level of task engagement and changes that are shared by both levels of task engagement. **c**. Two separate groups of participants underwent EEG-fMRI recording while listening to auditory tones. One group actively responded to the tones while the other group passively listened. **d**. The fMRI timeseries for 188 cortical and 2 thalamic ROIs were extracted, and the instantaneous phase coherence was calculated between each pair of ROIs at each timepoint. **e**. EEG data epochs were classified as alert or drowsy.

Within *Active* and *Passive* tasks separately, for each edge timecourse, we quantified the effect of alertness level predicting phase coherence (**Figure S1a**), and we identified edges that had sufficiently strong evidence in favor of this *State* model’s prediction using a Bayes Factor (BF) (i.e., *ln(BF) > 2*; see **Methods**) (**Figure S1b**). In the following comparisons, we focused on the topography of thalamocortical, frontoparietal, and auditory network edges in the *Drowsy>Alert* contrast as per pre-registered hypotheses. (See **Figure S2** for whole-brain results.)

### Low alertness is associated with increased thalamocortical synchronization during passive listening but not during active task engagement

With decreasing alertness, passive listening was associated with stronger and more widespread thalamocortical synchronization, in comparison to active task engagement. For both tasks, we observed increased phase coherence in the *Drowsy>Alert* contrast between bilateral thalami and specific cortical (i.e., Visual, Somatomotor, Dorsal Attention, Ventral Attention) networks (for [number of edges that passed threshold / total number of available edges] 61/216 edges; *Active* ln(BF): 7.34 ± 0.62; *Passive* ln(BF): 10.23 ± 0.76) (**Figure 2**: *Thalamocortical*, top row). These thalamocortical network edges included nodes distributed around the precentral and postcentral gyri (Somatomotor Network), supramarginal gyrus and posterior part of the cingulate gyrus (Ventral Attention Network), superior parietal lobule (Dorsal Attention Network), and occipital regions of the Visual Network. However, we also found additional widespread increased thalamocortical phase coherence that was specific to the *Passive* task in the *Drowsy>Alert* contrast across all cortical networks (for 107/376 edges; *Passive* ln(BF): 7.86 ± 0.43) (**Figure 2**: *Thalamocortical*, bottom row), although with higher distribution in parietal and occipital regions. These thalamocortical network edges with the highest BF came from the precentral and postcentral gyri (Somatomotor Network); bilateral inferior temporal gyrus, superior parietal lobule and lateral occipital cortex (Dorsal Attention Network); right precuneus and right frontal pole (Control Network); and throughout the occipital cortex (Visual Network). Thalamocortical increases in phase coherence that were shared by both tasks, as well as those specific to the *Passive* task, largely stemmed from the left thalamus (**Figure S3**: *Thalamocortical*).

**Figure 2:**
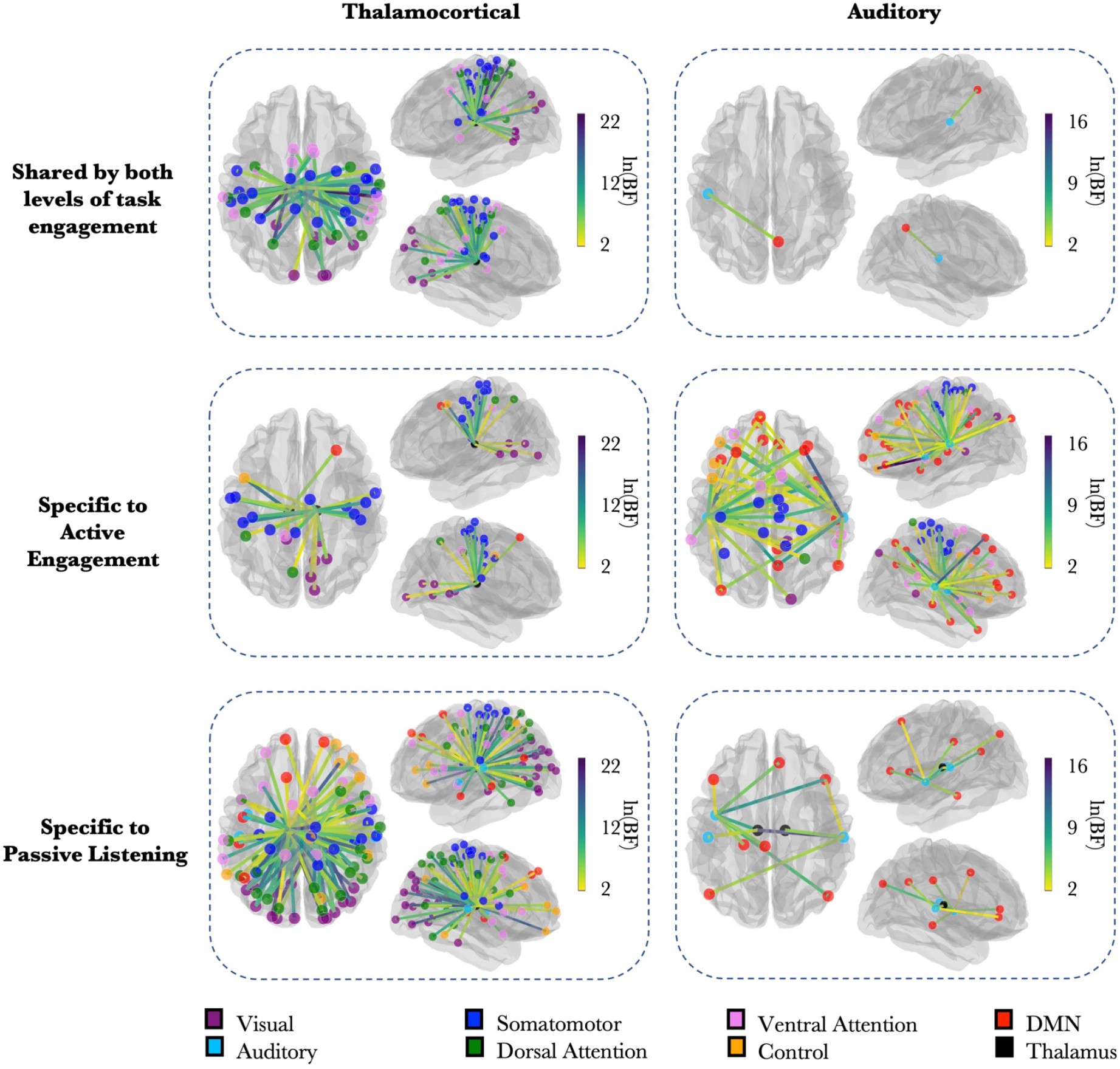
Spatial topography of thalamocortical and auditory network phase coherence. (Thalamocortical column) Glass brains show the thalamocortical network edges that had sufficiently strong evidence in favor of the *State* model for the *Drowsy>Alert* contrast (see **Methods**). Edge color denotes the BF. Node color denotes the ROI’s network. **(Auditory column)** Glass brains show the network edges with the Auditory module that had sufficiently strong evidence in favor of the *State* model for the *Drowsy>Alert* contrast. **(Top row)** Changes in phase coherence that were shared by both levels of task engagement. **(Middle row)** Phase coherence changes specific to the *Active* task. **(Bottom row)** Phase coherence changes specific to the *Passive* task.

We found few thalamocortical network edges that had increased phase coherence during the *Active*, but not *Passive*, task in the *Drowsy>Alert* contrast (for 26/368 edges; *Active* ln(BF): 6.53 ± 0.69) (**Figure 2**: *Thalamocortical*, middle row). These edges extended to the precentral and postcentral gyri of the Somatomotor Network, as well as the lingual gyrus of the Visual Network. They largely stemmed from the right thalamus (**Figure S3**: *Thalamocortical*).

### Low alertness is associated with increased auditory network synchronization during active task engagement but not during passive listening

We anticipated altered phase coherence with auditory brain regions during decreasing alertness because both tasks in this study involved auditory stimuli, and auditory-related functional reorganization has been observed between wakefulness and sleep stages ^23^. We found that with decreasing alertness, active task engagement, but not passive listening, was associated with widespread auditory network synchronization. This increased phase coherence in the *Drowsy>Alert* contrast during the *Active*, but not *Passive*, task spanned all cortical networks (for 56/736 edges; *Active* ln(BF): 4.78 ± 0.37) (**Figure 2**: *Auditory*, middle row). The auditory network edges with the strongest increase in phase coherence were largely distributed around the bilateral supplementary motor cortex (Somatomotor Network); left frontal pole (Control Network); right superior frontal gyrus and right precuneus (DMN); and right insula (Ventral Attention Network). This increased phase coherence with left and right auditory nodes was equally distributed across the ipsilateral and contralateral hemispheres (**Figure S3**: *Auditory*).

In comparison, we found minimal auditory network synchronization during decreasing alertness that was specific to passive listening. These auditory network edges that had increased phase coherence in the *Drowsy>Alert* contrast during the *Passive*, but not *Active* task, were distributed in the DMN and bilateral thalami (for 12/192 edges; *Passive* ln(BF): 5.36 ± 0.90) (**Figure 2**: *Auditory*, bottom row). DMN nodes included the right anterior cingulate cortex, right inferior gyrus, and bilateral angular gyrus.

There was only one auditory network edge shared by both *Active* and *Passive* tasks that had increased phase coherence in the *Drowsy>Alert* contrast (1/184 edges; *Active* ln(BF): 2.573; *Passive* ln(BF): 5.892) (**Figure 2**: *Auditory*, top row). This edge was shared by a left auditory node and the right precuneus from the DMN.

### Low alertness is associated with frontoparietal synchronization during active task engagement but predominantly parietal synchronization during passive listening

Finally, we explored the spatial topography of frontoparietal synchronization during decreasing alertness because we expected increased frontoparietal synchronization (i.e., among Dorsal Attention, Ventral Attention, and Control Networks) to reflect the interaction between decreasing alertness and sustained decision-making ^24–26^. We found that active task engagement, but not passive listening, was associated with long-range frontoparietal synchronization during decreasing alertness. These long-range frontoparietal network edges that had increased phase coherence in the *Drowsy>Alert* contrast during the *Active* but not *Passive* task, extended from medial, frontopolar, and dorsolateral prefrontal areas toward bilateral posterior parietal hubs (82 edges; ln(BF): 5.565 ± 0.448) (**Figure 3**: middle row).

**Figure 3:**
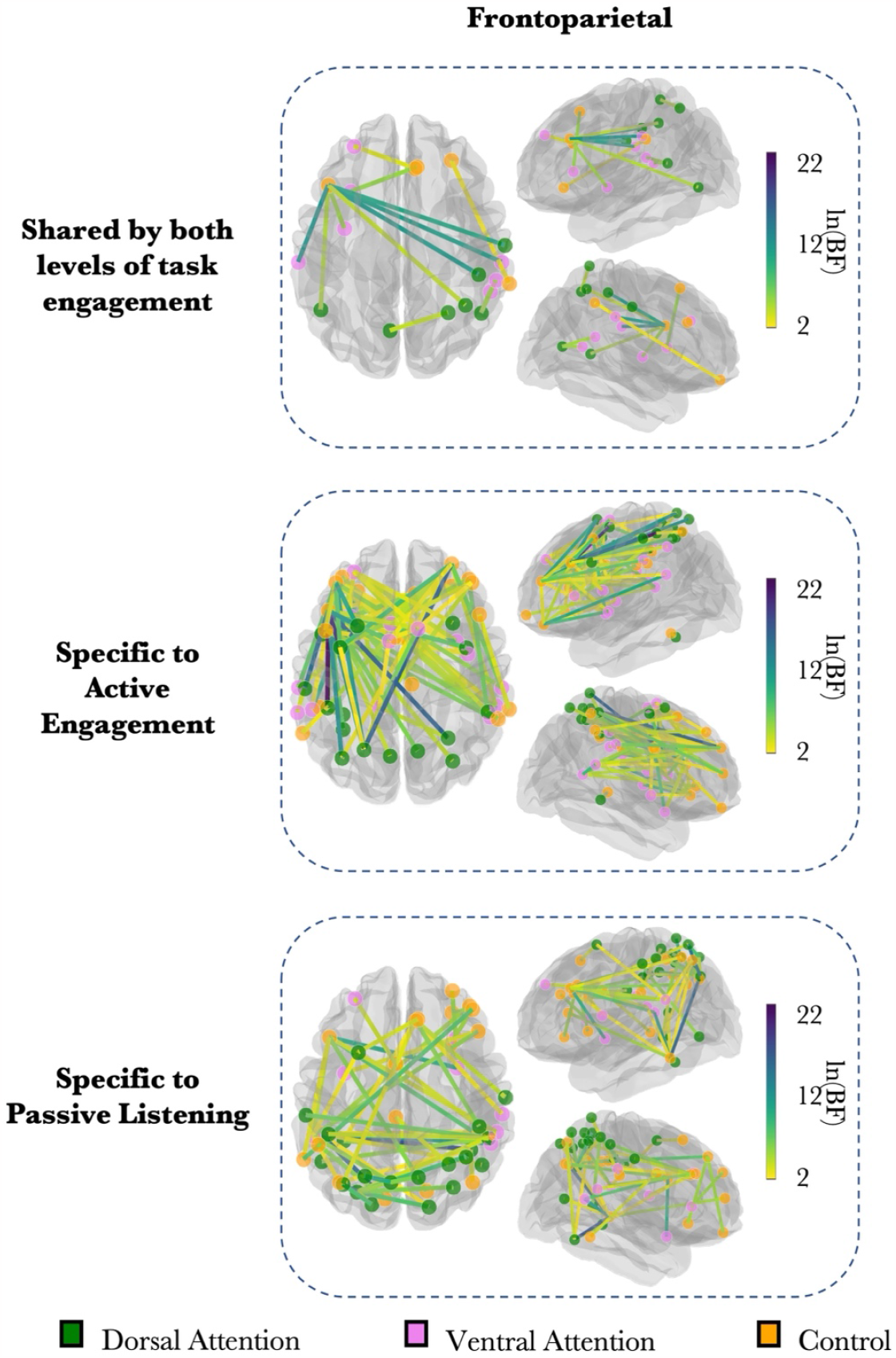
Spatial topography of frontoparietal network phase coherence. Glass brains show the frontoparietal network edges (within the Dorsal Attention, Ventral Attention, and Control Networks) that had sufficiently strong evidence in favor of the *State* model for the *Drowsy>Alert* contrast. Edge color denotes the BF. Node color denotes the ROI’s network. **(Top row)** Changes in phase coherence that were shared by both levels of task engagement. **(Middle row)** Phase coherence changes specific to the *Active* task. **(Bottom row)** Phase coherence changes specific to the *Passive* task.

In comparison, decreasing alertness during passive listening was associated with more parietal synchronization. We found fewer frontoparietal network edges and more posterior parietal edges that had increased phase coherence in the *Drowsy>Alert* contrast during the *Passive*, but not *Active*, task. This increased phase coherence included a heavily connected interhemispheric network of posterior parietal and temporoparietal edges and little frontal involvement (48 edges; ln(BF): 5.491 ± 0.49) (**Figure 3**: bottom row).

We found that with decreasing alertness, little frontoparietal synchronization was common to both levels of task engagement. That is, *Active* and *Passive* tasks only shared a few common frontoparietal network edges that had increased phase coherence in the *Drowsy>Alert* contrast (13 edges; *Active* ln(BF): 7.544 ± 1.625; *Passive* ln(BF): 5.025 ± 0.568) (**Figure 3**: top row).

## DISCUSSION

Here, we showed how thalamocortical, auditory, and frontoparietal dynamics change during transitions of consciousness across different levels of task engagement. We found stronger long-range frontoparietal and auditory network synchronization with decreasing alertness when participants were actively engaged in an auditory task, but not when passively listening to auditory tones. With decreasing alertness, passive listening, but not active task engagement, was associated with increased synchronization within the parietal lobes and widespread increased thalamocortical synchronization. Alertness fluctuations and cognitive processing are associated with overlapping brain dynamics, and an understanding of how these two processes jointly modulate their overlapping brain circuits is needed to advance towards an integrated framework of consciousness and cognition ^18^.

It has been proposed that **thalamocortical** networks herald the early process of falling asleep ^9^ and losing consciousness during sedation ^27^, with the thalamus leading this transition ^28^. The dramatic changes in thalamocortical networks for the *Passive* task support this prediction. Further, it has been proposed that brain networks that are active during a cognitive task may remain active even if the participant is fluctuating in alertness ^25,29^, a hypothesis that is also supported by the little change in thalamocortical phase coherence when alertness decreased during the *Active* task.

It has been long proposed that large-scale **frontoparietal organization** underlies the integration of cognitive processes and cross-network interaction ^30–33^. The brain’s flexibility to functionally reorganize during decreasing alertness according to cognitive task load can be interpreted under a cognitive framework of neural information integration ^34^. We assume that spatiotemporal information integration changes with different levels of alertness for the same task, and phase coherence can reflect the brain network resources required to sufficiently integrate information. We’ve recently proposed that this integration separately depends on both alertness level and task ^35^, the increased frontoparietal synchronization observed in the *Drowsy>Alert* contrast during the *Active*, but not *Passive* task, suggests that during low alertness additional cognitive resources are required to achieve the level of information integration that is associated with high alertness. These frontoparietal network edges did not show increased phase coherence in the *Drowsy>Alert* contrast during the *Passive* task, suggesting that they contribute specifically to task engagement in the drowsy state and are not driven by sleep onset alone. Our results support the notion that the brain operates flexibly and has available resources to recruit when challenged ^36^.

**Auditory** processing is largely preserved in sleep ^29,37^, but it was recently shown that it involves a functional reorganization across wakefulness and sleep stages ^23^. Our work suggests that this reorganization begins as early as the transition from high to low alertness and is dependent on task engagement. Increased phase coherence with auditory regions during decreasing alertness was largely specific to the *Active* task, particularly among prefrontal brain areas and the Somatomotor Network. The *Passive* task, in comparison, was associated with little change in phase coherence and only showed sparse increases in synchronization with the DMN and bilateral thalami. In the *Active* task, participants were not only hearing tones, but also making a decision to press one of two buttons, thus maintaining perceptual decision-making processes. It is therefore task engagement that drives auditory integration when alertness fades.

Throughout the day, alertness fluctuations depend on a multiplicity of factors ^38^. In a third of the human population, low alertness can easily be triggered by repetitive tasks or passive task engagement that enables mind-wandering ^39,40^. In the current study, we took advantage of the natural tendency of our participants to fluctuate between high and low alertness, and labeled their states as alert or drowsy based on robust computational methods applied to the EEG signal dynamics ^22^. We used fMRI BOLD signal phase coherence to characterize the brain as a complex system with non-linear interactions between wakefulness and cognitive task engagement ^41^. There is an established view of the dynamics across different levels of task engagement, along with the anticorrelations observed among large brain networks ^42,43^, which needs to be integrated with the literature on wakefulness versus unconsciousness for subcortical and cortical network differences ^44^. In this study, we offer the same cognitive neuroscience framework for both, conceptually and analytically. While apparently overlapping frontoparietal networks have been systematically implicated in executive function, decision-making, and cognitive control tasks ^16^, the same networks have been signaled as key for decreased consciousness states such as sleep, anesthesia, vegetative state, and minimally conscious state ^17^. This apparent field split between cognition and consciousness ^45^ is directly put to the test in the current experiments, showing a nuanced interaction between cognition, engagement, and transitions to lower consciousness states.

Cautious interpretation should be made on indirect neural measures. This work relied on fMRI phase coherence as a measure of dynamic brain regional functional coordination. We favored phase coherence over functional connectivity as it is an instantaneous measure, and alertness fluctuations vary on the timescale of seconds ^22^. Phase coherence has also been used successfully in other recent work ^21,46–49^.

The brain network dynamics identified in active task engagement during low alertness may not be sufficient to maintain cognition in the same way that it is maintained during high alertness ^22,24^. While some functional brain configurations may allow us to perform optimally when drowsy and in other impaired states of consciousness, others may be maladaptive and correspond with poor performance and pathological mechanisms. Therefore, while our results may infer a causal linkage between behavior and different brain network dynamics modulated by alertness, this causality requires further exploration ^50^.

## METHODS

This work was part of a larger preregistered project that can be viewed here: https://osf.io/nua4y.

### Data acquisition

The *Active* task study was conducted as described in a previously published only-EEG study ^51^, except that the current paradigm involved a simultaneous EEG-fMRI set-up. Data collection took place in 2013. The *Passive* task study also involved a simultaneous EEG-fMRI set-up, and data collection took place in 2017.

#### Participants

In the *Active* task, 25 right-handed healthy volunteers (12 females; age 28 ± 4 years old) with normal hearing and no neurological or psychiatric conditions participated in the study. In data analysis, one subject was excluded due to a corrupted EEG data file, two subjects were excluded due to high motion throughout the scan, and two subjects were excluded due to artifacts in the EEG data that could not be removed sufficiently for the drowsiness algorithm to properly work. The *Passive* task had 26 volunteers (14 females; age 25.15 ± 5.28 years old) with the same screening criteria as in the *Active* task. One subject was excluded due to high motion throughout the scan, two subjects were excluded due to artifacts in the EEG data that could not be removed sufficiently for the drowsiness algorithm to properly work, and three subjects were excluded because at least ⅚ of their available runs contained NREM stage 2 sleep. In the end, 20 *Active* task subjects and 20 *Passive* task subjects were used in all analyses. Participants were recruited using the Cambridge Psychology SONA system, provided informed consent prior to the study, and received a compensation of £10 per hour for participation. The protocols were approved by the Cambridge Psychological Research Ethics Committee before any testing began.

#### Stimuli

*Active* task participants were tested lying down in the MRI scanner while wearing in-ear etymotics, which presented harmonic complex tones that differed in their perceived location (ranging 0°-60° left and right of midline). Tones were recorded in a free field prior to data collection using in-ear microphones. Participants were asked to indicate whether the tone originated from right or left of their midline by pushing a right-or left-side button on a response box with the right or left thumb correspondingly.

The *Passive* task was divided into two sessions: awake and drowsy/sleep. Each of the two sessions consisted of three runs. In the first run, participants had five minutes of resting state followed by five minutes of an auditory event-related paradigm including three pure tones (250, 1000, 4000 Hz). The last two runs consisted of either, in randomized order, 2) only ascending or 3) only descending tones (14 minutes each). There were 11 pure tone stimuli (125, 177, 250, 354, 500, 707, 1000, 1414, 2000, 2828, 4000 Hz) that were presented either in ascending or in descending order. Accordingly, pure tone bursts of either the lowest or highest frequency were presented for two seconds (equivalent to the length of 1 TR) before moving on to the next higher or lower frequency until all 11 tones were played. One sequence of tones was then followed by a pause of four seconds, and this procedure was repeated 30 times per run. In total, each participant underwent two ascending and two descending runs.

#### Procedure

In preparation for the experiment, participants were instructed to refrain from drinking alcohol the night before and coffee on the day of the experiment. Before entering the MRI scanner, participants were fitted with an EGI electrolyte 128-channel cap developed by Electrical Geodesics, as well as earphones for stimulus presentation.

In the *Active* task, participants were asked to have their eyes closed and respond as quickly and accurately as possible to the stimuli presented. In each stimulus trial, 5-8 seconds passed before the presentation of the tone, and the next trial began if a response was not detected within five seconds. Participants were encouraged to relax and not worry about falling asleep as the tone volume would increase and wake them up if needed. They were allowed about five minutes to relax and become sleepy, after which the session commenced. The entire task lasted approximately 50-60 minutes.

*Passive* task participants were instructed to keep their eyes closed and stay as still as possible. They were requested to remain awake during the awake session and encouraged to fall asleep during the drowsy/sleep session. Tones were presented during both these alert and sleep sessions, and the sessions combined lasted about an hour.

#### Scanning acquisition parameters

The *Active* task was conducted using the Siemens 3T Tim Trio MRI scanner (32-channel head coil), while the *Passive* task was conducted using the Siemens 3T Prisma (20-channel head coil), both at the Medical Research Council Cognition and Brain Sciences Unit (MRC CBU) at the University of Cambridge. EEG data were collected using a Mac computer and the EGI Netstation software (www.egi.com).

In both tasks, T2*-weighted images were acquired with an echo-planar imaging (EPI) sequence using axial slice orientation (slice thickness: 3.75 mm; repetition time: 2000 ms; echo time: 30 ms; flip angle: 78°; field of view: 192 x 192 mm; voxel size: 3 x 3 x 3.75 mm). A T1-weighted anatomical image was also acquired (repetition time: 2250 ms; echo time: 2.98 ms; flip angle: 9°; voxel size: 1 x 1 x1 mm).

### Data processing

#### EEG preprocessing

EEG data were processed using the EEGLAB toolbox ^52^ with Matlab 2019b (MathWorks, Inc., MA, USA). EEG data acquired during fMRI scanning are subject to magnetic field gradient and ballistocardiographic (BCG) artifacts. We addressed these two artifacts using the EEGLAB plug-in provided by the University of Oxford Centre for Functional MRI of the Brain (FMRIB) ^53^. We also performed a manual artifact rejection of the EEG data and inputted clean data segments into an independent component analysis (ICA) to check for and remove any noise-related artifacts that still remained. EEG data were downsampled to 250 Hz, filtered between 1-30 Hz, and epoched. Missing and bad channels were interpolated using spherical spline interpolation.

#### EEG-defined drowsiness

To measure alertness at an electrophysiological level, EEG data from both *Active* and *Passive* tasks were inputted into an automatic algorithm for drowsiness classification, in which each four-second data epoch was classified as “alert” or “drowsy” ^22^. The algorithm is loosely based on the Hori system that breaks down the wake-sleep transition into nine stages ^5^. The algorithm uses whole-brain information from EEG coherence and the variance explained by the low-frequency spectral power bands to classify epochs in a subject-specific way.

#### MRI preprocessing

MRI data were preprocessed using the Analysis of Functional NeuroImages (AFNI) toolbox ^54^ and FreeSurfer (http://surfer.nmr.mgh.harvard.edu/). We extracted masks for ventricles, gray matter, and white matter from the anatomical image. We removed the first 10 volumes of each subject’s functional data. fMRI preprocessing steps included slice-time correction, motion correction, coregistration, normalization of the functional and structural data into standard space, spatial smoothing (4-mm full width at half maximum), and temporal filtering (0.01-0.1 Hz). A nuisance regression was performed using motion parameters, as well as the first 3 principal components from the ventricles and white matter ^55,56^. We identified voxels that had low temporal variance (standard deviation < 3) to compute a group-level full-brain mask for subsequent ROI extraction.

#### fMRI analysis

fMRI ROI timeseries extraction and visualization were performed in Python v.3.8 (Python Software Foundation, http://python.org), and Nilearn (http://nilearn.github.io). We used a whole-brain cortical parcellation that divides the brain into seven networks and 200 parcels ^19,57^. Voxel time courses were extracted for each parcel and spatially averaged to calculate the timeseries of a single ROI. We additionally extracted two timeseries for the left and right thalamus using the AAL atlas ^20^. The group-level mask had poor coverage in the frontal and temporal poles, coinciding with the Schaefer/Yeo Limbic Network. These 12 Limbic ROIs were consequently removed from analyses, so there were 188 cortical + 2 thalami = 190 ROIs total. Because both tasks involved auditory stimuli, we extracted 4 ROIs (*17Networks_LH_SomMotB_Aud1 / Aud2, 17Networks_RH_SomMotB_Aud1 / Aud2*) from the Schaefer/Yeo Somatomotor Network to create an Auditory module.

Our method to calculate fMRI phase coherence has been previously reported ^21^. The result was a 190 x 190 phase coherence matrix at each timepoint that contained a phase coherence value for every pair of ROIs.

#### Data selection criteria

We applied several data selection criteria to ensure that data from the two tasks were comparable. In the *Active* task, participants’ continuous behavioral responses confirmed that they had not entered NREM stage 2 sleep, which is characterized by unresponsiveness ^58^. The *Passive* task had six runs per subject, and participants were susceptible to falling into NREM light sleep. For the statistical analyses, we selected *Passive* task runs that only comprised wake and/or NREM1 according to manual sleep staging. We delayed the fMRI time series by 3 TRs (6 seconds) to approximate one-to-one mapping between electrophysiologically defined drowsiness and fMRI functional coordination because it has been estimated that the BOLD signal follows neural processes with a 4-to 6-second delay ^59,60^. With the fMRI data, a Euclidean norm was calculated from the six motion parameters to get a single metric of motion at each TR. We excluded each high-motion volume (> .25mm), as well as one TR before and after. Finally, we selected data points that remained in the same EEG drowsiness state for both the current and subsequent TR. (This step helped focus our analyses on canonical alert and drowsy states.) This final subset of the data was used for all statistical analyses. Following these data selection criteria, the *Active* task had 7031 total samples assigned as alert (351.55 ± 126.35 samples per subject) and 6931 total samples assigned as drowsy (346.55 ± 174.79 samples per subject), which were inputted into the statistical analyses. The *Passive* task had 8568 total samples assigned as alert (428.40 ± 213.38 samples per subject) and 10433 total samples assigned as drowsy (521.65 ± 244.93 samples per subject), which were inputted into the statistical analyses.

#### Statistical analysis

Statistical analyses were performed in R v.4.0, using the libraries lmerTest ^61^ and bayestestR ^62^. For each unique pair of ROIs (i.e., each ROI edge), we quantified the effect of State (drowsy versus alert) predicting phase coherence in a linear mixed-effects model, with subject as random effect: *phase_coherence ∼ State + (1*|*subject)*. Our null model only included subject as random effect (i.e., *phase_coherence ∼ (1*|*subject)*). We used the natural log (*ln*) of the BF to conservatively constrain the results to ROI edges that had sufficiently strong evidence in favor of the *State* model’s prediction (i.e., *ln(BF*_*State model*_*) > 2*, roughly equivalent to *BF*_*State model*_ *> 7*, when *ln(BF*_*null*_*) = 0*) ^63,64^. A BF is used to compare two statistical models against each other, and it quantifies the likelihood that one model (i.e., *State* model) describes the data better than the other model (i.e., null model). A higher BF indicates greater support in favor of the *State* model ^65^.

The diagram summary of the results (**Figure 1a**) was created with BioRender.com.

Analyses were conducted using computational resources from the NIH HPC Biowulf cluster (http://hpc.nih.gov).

## Acknowledgments

This work was supported by the NIH OxCam Scholars Program (to S.K.), The Wellcome Trust (WT093811MA to T.A.B. and UNS49938 to A.A.).

