## Supplementary Material for "Brain network dynamics in transitions of consciousness reorganize according to task engagement"

Acknowledgments
This work was supported by the NIH OxCam Scholars Program (to S.K.), The Wellcome Trust (WT093811MA to T.A.B. and UNS49938 to A.A.).

Declaration of interests
The authors declare no competing interests.

### SUPPLEMENTARY FIGURES

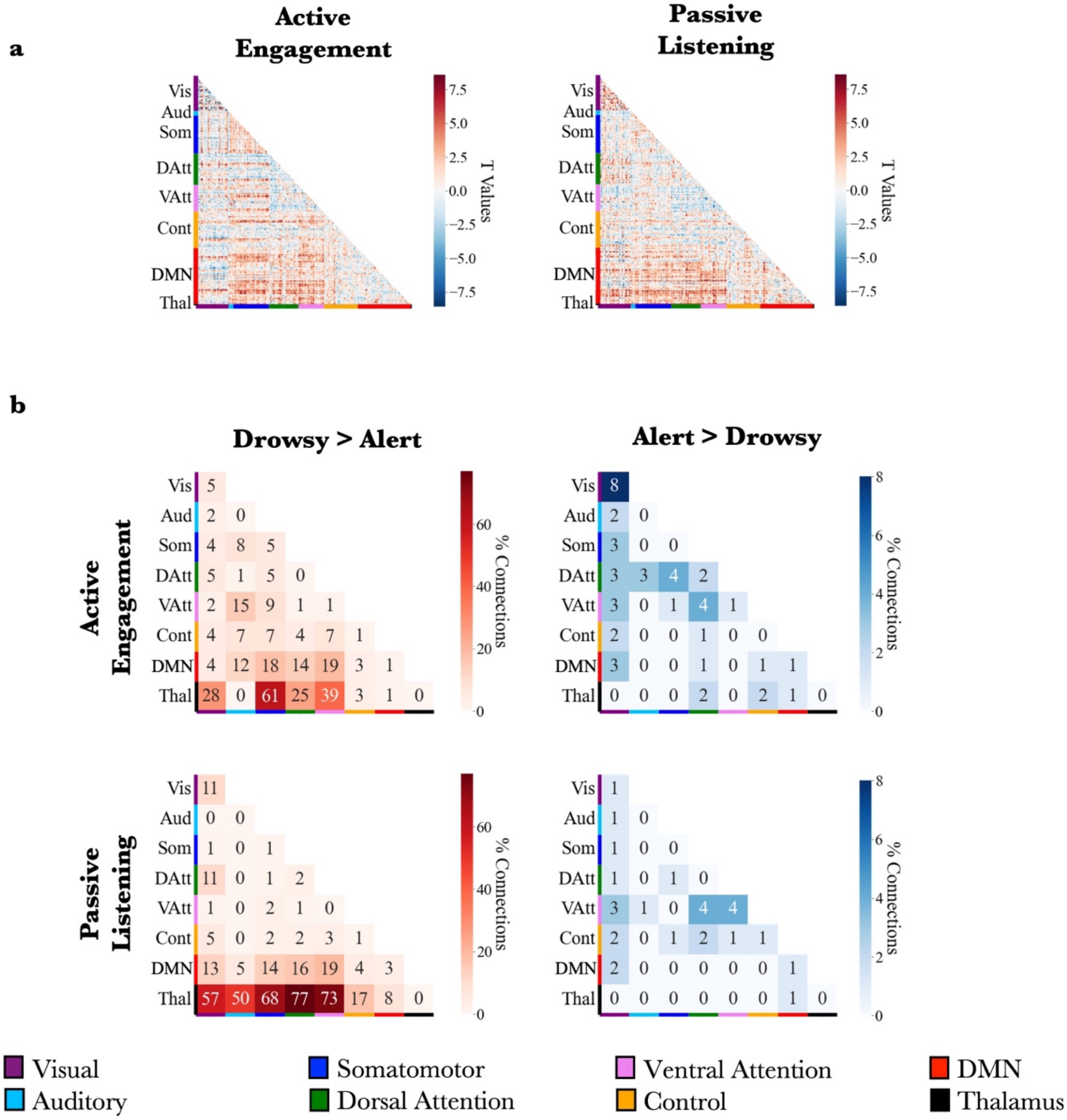


**Supplementary Figure 1: Alertness level predicts phase coherence. a.** The effect of alertness level predicting phase coherence for each ROI edge. Higher t-value denotes *Drowsy>Alert*. Low t-value denotes *Alert>Drowsy*. **b.** Each square in each matrix shows the percentage of ROI edges between two networks that had sufficiently strong evidence in favor of the test model's prediction (i.e., *ln(BF) > 2*) (see **Methods**).

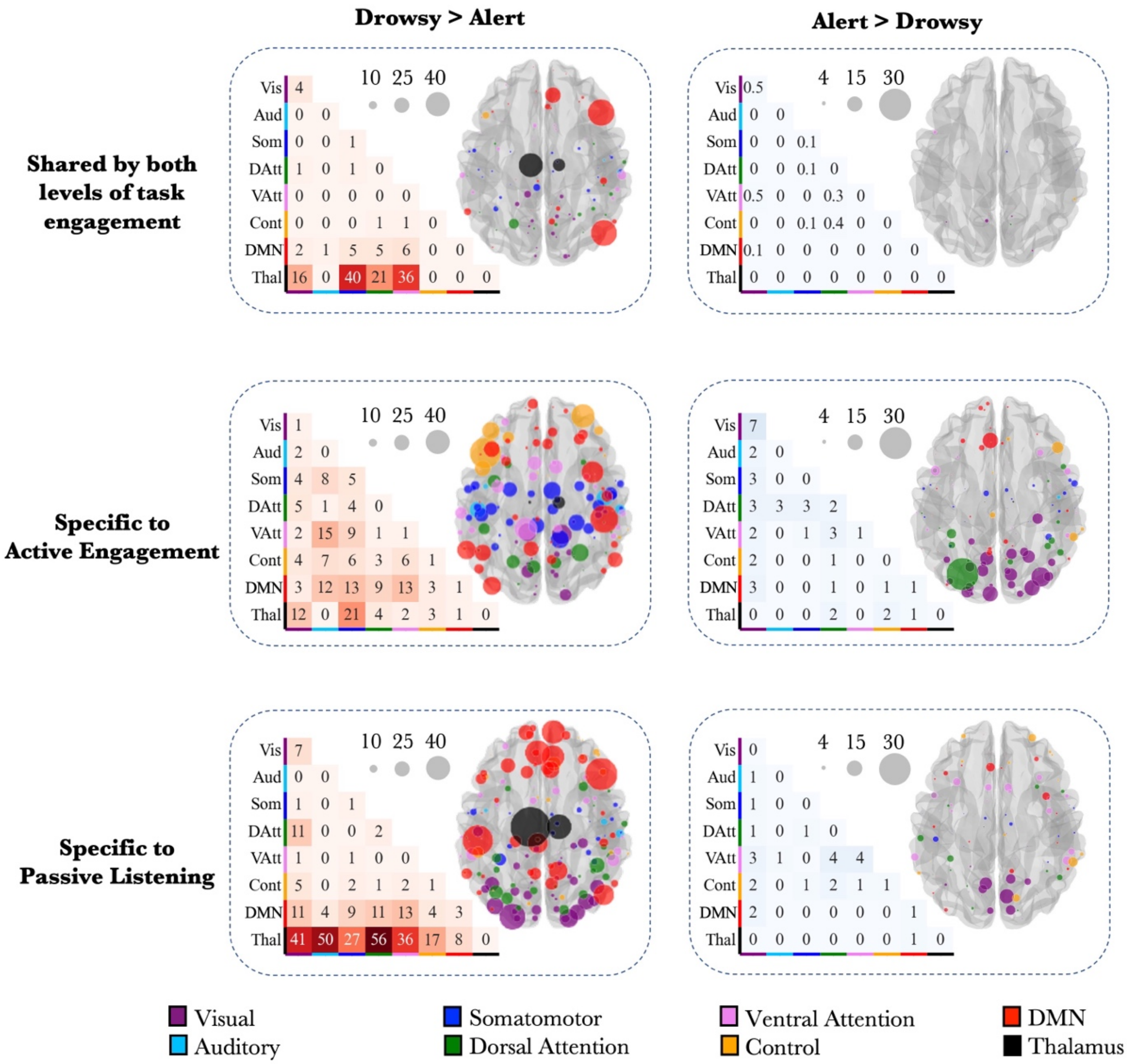


**Supplementary Figure 2: Whole-brain changes in phase coherence.** Each square in the matrix denotes the percentage of edges between two networks that had sufficiently strong evidence in favor of the test model's prediction. Each glass brain shows the number of edges that had sufficiently strong evidence in favor of the test model, for each ROI node. A larger node in the glass brain denotes a higher number of edges connecting with that node. **(Top row)** Changes in phase coherence that were shared by both levels of task engagement. **(Middle row)** Phase coherence changes specific to *Active* task. **(Bottom row)** Phase coherence changes specific to *Passive* task.

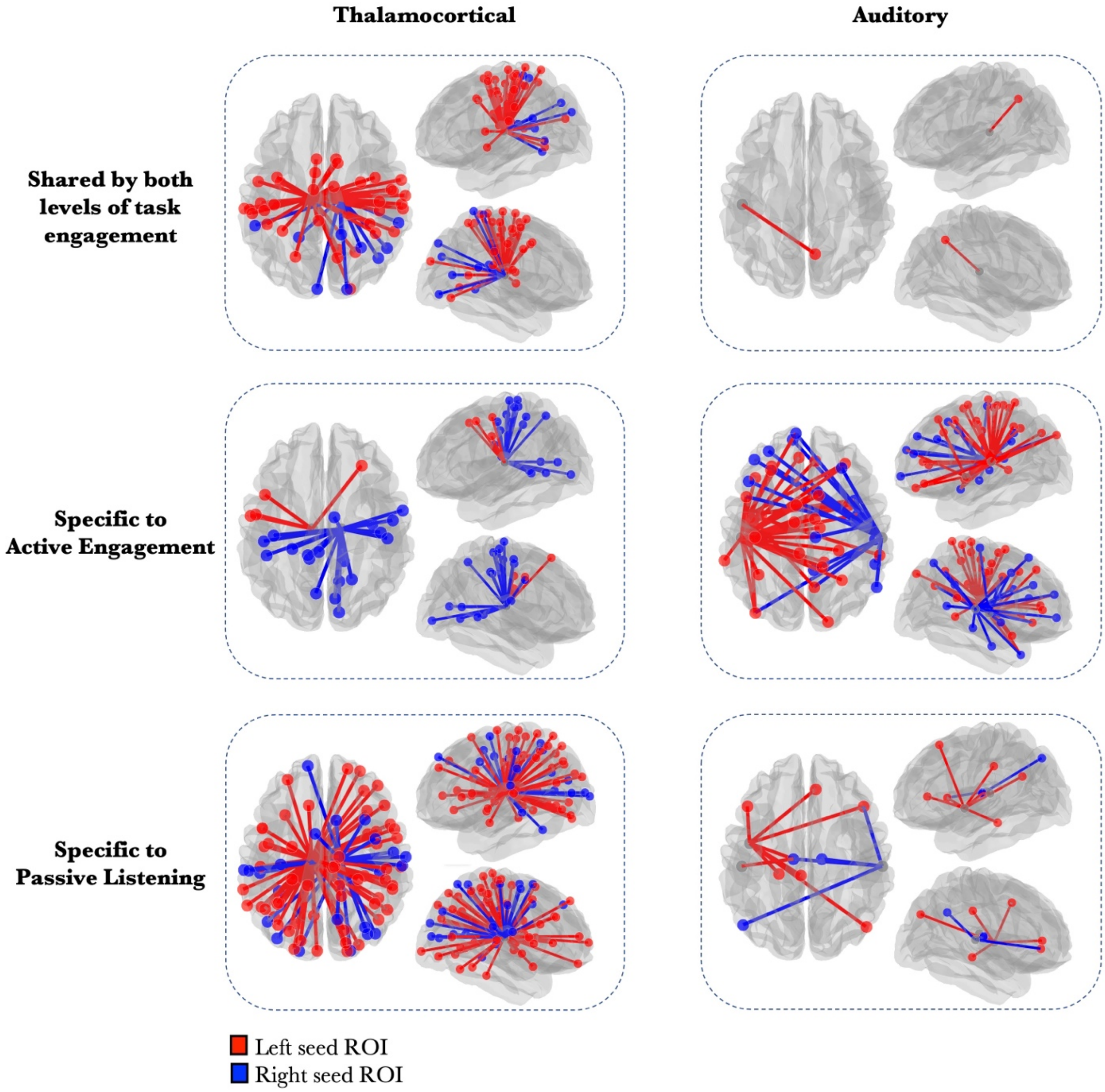


**Supplementary Figure 3: Laterality of thalamocortical and auditory network phase coherence. (Thalamocortical column)** Glass brains show the thalamocortical network edges that had sufficiently strong evidence in favor of the test model for the *Drowsy>Alert* contrast. Edge and node color denote whether the edge stems from the right or left thalamus. **(Auditory column)** Glass brains show the network edges with the Auditory module that had sufficiently strong evidence in favor of the test model for the *Drowsy>Alert* contrast. Edge and node color denote whether the edge stems from the right or left auditory nodes. **(Top row)** Changes in phase coherence that were shared by both levels of task engagement. **(Middle row)** Phase coherence changes specific to *Active* task. **(Bottom row)** Phase coherence changes specific to *Passive* task. Thalamocortical edges shared by both levels of task engagement: 40 edges with left node, 21 edges with right node. Thalamocortical edges specific to Active Engagement: 5 edges with left node, 21 edges with right node. Thalamocortical edges specific to Passive Listening: 66 edges with left node, 41 edges with right node. Auditory edges shared by both levels of task engagement: 1 edge with left node, 0 edges with right node. Auditory edges specific to Active Engagement: 36 edges with left node, 20 edges with right node. Auditory edges specific to Passive Listening: 8 edges with left node, 4 edges with right node.
